# BleTIES: Annotation of natural genome editing in ciliates using long read sequencing

**DOI:** 10.1101/2021.05.18.444610

**Authors:** Brandon K. B. Seah, Estienne C. Swart

**Affiliations:** Max Planck Institute for Developmental Biology, Max-Planck-Ring 5, 72076 Tübingen, Germany

## Abstract

**Summary:** Ciliates are single-celled eukaryotes that eliminate specific, interspersed DNA sequences (internally eliminated sequences, IESs) from their genomes during development. These are challenging to annotate and assemble because IES-containing sequences are much less abundant in the cell than those without, and IES sequences themselves often contain repetitive and low-complexity sequences. Long read sequencing technologies from Pacific Biosciences and Oxford Nanopore have the potential to reconstruct longer IESs than has been possible with short reads, and also the ability to detect correlations of neighboring element elimination. Here we present BleTIES, a software toolkit for detecting, assembling, and analyzing IESs using mapped long reads.

**Availability and implementation:** BleTIES is implemented in Python 3. Source code is available at https://github.com/Swart-lab/bleties (MIT license), and also distributed via Bioconda.

**Contact:** Contact

**Supplementary information:** Benchmarking of BleTIES with published sequence data.

## Text

### Introduction

Ciliate cells have two different types of nuclei: somatic macronuclei (MAC) and germline micronuclei (MIC). MACs develop from MICs during conjugation following meiosis and fertilization. During development, the genome is extensively edited, to excise thousands of interspersed, internally eliminated sequences (IESs) through an RNA-guided mechanism, accompanied by chromosome fragmentation, and segmental rearrangement, with significant variation in the nature and degree of editing between species (reviewed in: (Chalker *et al.*, 2013)).

The task of identifying such edits from sequence data is a specialized subset of structural variant detection, whereby characteristics peculiar to ciliates, i.e. short tandem repeats called ‘pointers’ (Chen *et al.*, 2014) or TA repeats characteristic of domesticated excisases (Klobutcher and Herrick, 1995) at IES junctions, can be exploited for annotation. When mapped against a MAC reference genome, genomic DNA reads that originate from MICs or developing MACs contain IESs or arrangements not present in the mature MAC. These are observed as inserts or clips in the aligned reads, and can be used to annotate IES and genome scrambling junctions and to reconstruct the IES sequences. Accurate IES annotations are necessary to analyze the effects of knockdown of candidate genome rearrangement genes by measuring IES retention across known loci, expressed as an “IES retention score” (Swart *et al.*, 2014; Denby Wilkes *et al.*, 2016). Software for annotating IESs, genome scrambling, and scoring retention have been developed for short read sequencing data, e.g. ParTIES (Denby Wilkes *et al.*, 2016), ADFinder (Zheng *et al.*, 2020), and SIGAR (Feng *et al.*, 2020).

Single-molecule long read sequencing from Pacific Biosciences (PacBio) and Oxford Nanopore offer advantages over short reads for IES prediction: (i) long reads several kbp long may span multiple IESs, unlike short reads (~300-400 bp insert size) which span either one or zero IES sites; (ii) the entire IES sequence can potentially be covered by the read; (iii) IESs containing repetitive sequences are more likely to be detected with more sequence context. Because each long read represents a single genomic DNA molecule, the correlation of IES excision or retention between neighboring IESs can be investigated. Existing software packages like ParTIES were designed for short reads, make assumptions inapplicable to long reads with higher error rates, and would not be easily adapted to accommodate them. Therefore, we developed BleTIES (Basic Long-read Enabled Toolkit for Interspersed DNA Elimination Studies) for IES detection and analysis from long read libraries.

### Description

BleTIES has a modular design: the main module for *de novo* IES reconstruction, MILRAA, depends on standard bioinformatics libraries Biopython, pysam, and htslib (Cock *et al.*, 2009; Li *et al.*, 2009). Input data are the MAC genome assembly (Fasta format) and genomic long reads mapped to it (BAM format (Li *et al.*, 2009)). If a read contains IESs, the mapper should report them as inserts relative to the assembly. PacBio circular consensus sequences (CCSs) are sufficiently accurate to infer IES coordinates directly, but naive calling of insert coordinates and sequences from uncorrected long reads (PacBio subreads or Nanopore reads) is inaccurate because of their high error rates. Therefore, with such reads MILRAA identifies clusters of inserts within a specified threshold distance, and if a cluster is above the minimum coverage cutoff, the insert and flanking 100 bp regions are extracted from each mapped read, and assembled to a more accurate consensus sequence with SPOA *(Vaser et al., 2017)*. The consensus is re-aligned to the reference, with the included flanking sequence helping to “anchor” the consensus in the reference, simultaneously predicting a more accurate insert position and sequence (Figure 1). Tandem repeats (pointers) and/or TA motifs flanking the insert are also annotated if present. The IES retention score is also reported.

**Figure 1.**
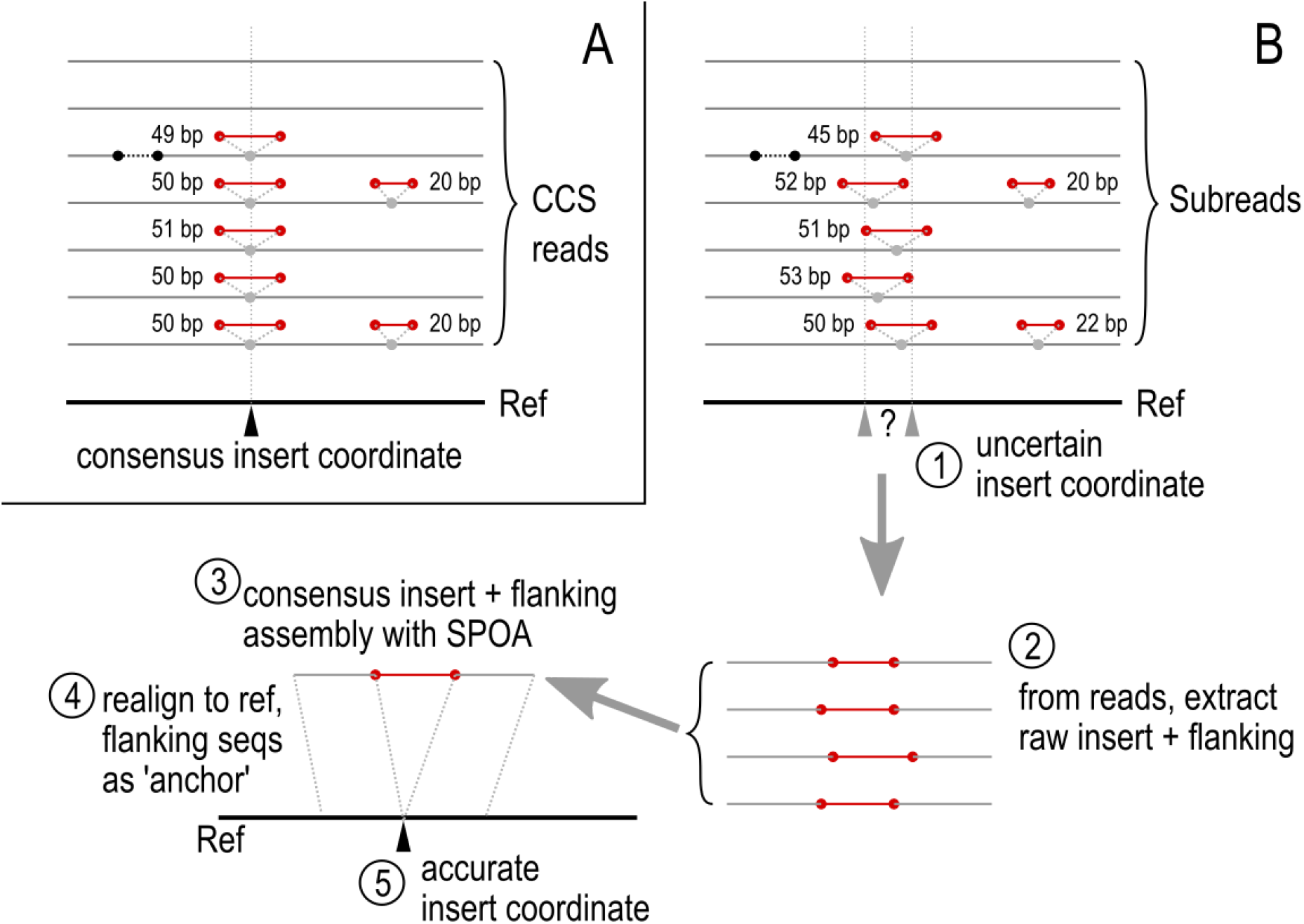
Overview of BleTIES MILRAA method to reconstruct IES junctions from error-corrected CCS reads vs. from subreads. (A) CCS reads have accuracy >99%, so the consensus insert coordinate from read mapping is adequate if coverage is sufficient. (B) (1) Subreads have higher error rates, so mapping alone is insufficient to define the insert position. (2) Therefore, clusters of adjacent inserts are identified, from which the insert sequence plus flanking +/- 100 bp are extracted from the subreads. (3) Extracted sequences are assembled with SPOA (Vaser *et al*., 2017). (4, 5) This consensus is realigned to the reference to obtain a more accurate estimate of the insert position.

In addition to the standard, per-IES retention score, BleTIES reports a per-read retention score: the fraction of annotated IES sites spanned by a read where the IES is not excised. This metric can be used to bin reads that have high IES retention, using the MILCOR module, useful for partial assembly of the MIC genome. MILTEL predicts alternative chromosome breakage sites by looking for reads bearing telomeric repeats in their clipped regions.

To test MILRAA against defined inputs and to estimate the coverage required for IES prediction, we simulated long read libraries containing mixtures of published MAC-only and MAC+IES sequences with pbsim2 v2.0.1 (Ono *et al.*, 2020) from *Paramecium tetraurelia* (Arnaiz *et al.*, 2012), whose predicted IESs are mostly <100 bp in length. We found MAC+IES coverage of 20× with PacBio subreads was adequate for identifying 93% of the IES junctions, almost all of which had flanking TA sequences reconstructed; about 48% of original IESs were assembled exactly, and 70% with a maximum of 1 mismatch and 1 indel. As expected, more IESs were identified and correctly assembled with higher MAC+IES coverage (Supplementary Information).

MILRAA was also tested with real mixed libraries of MIC and MAC (NCBI accession: SAMN12736122) mapped against a MAC genome reference (Sheng *et al.*, 2020) for *Tetrahymena tetraurelia*, with 7551 previously characterized IESs (median length 2.78 kbp (Hamilton *et al.*, 2016)). Two-thirds of these could be recovered (median length ~2 kbp) with median sequence identity ~99%, at ca. 50-60× read coverage. In addition, about 2800 new IESs were predicted. IESs that were not recovered were on average longer (median ~5 kbp), hence insufficient reads could span the entire IES for prediction by MILRAA (Supplementary Information). These could be resolved by binning out IES-containing reads with MILCOR and reassembling them, or by increasing the average read length.

### Conclusion

We have demonstrated the benefits of single-molecule long read sequencing data over short reads in annotating genome editing in ciliates, using our software tool BleTIES. These include the detection and assembly of longer IESs, and direct reporting of IES sequences and retention scores. One SMRT cell of the PacBio Sequel II platform currently (2021) yields ~300 Gbp of subreads, and ~20 Gbp of error-corrected CCS reads. Given modest ciliate genomes, e.g. 72 Mbp (MAC) and 98 Mbp (MIC) for *P. tetraurelia* (Guérin *et al.*, 2017), several IES retention experiments could potentially be analyzed with a single multiplexed sequencing run.

BleTIES is written in Python 3 and released under the MIT license; the version described here is v0.1.9. Source code and documentation are available at https://github.com/Swart-lab/bleties (archived at Zenodo doi:10.5281/zenodo.4723565). The package is also distributed via Bioconda (Grüning *et al.*, 2018) under the recipe ‘bleties’.

## References

## Bibliography

Arnaiz,O. et al. (2012) The Paramecium germline genome provides a niche for intragenic parasitic DNA: evolutionary dynamics of internal eliminated sequences. PLoS Genet., 8, e1002984.

Chalker,D.L. et al. (2013) Epigenetics of ciliates. Cold Spring Harb. Perspect. Biol., 5, a017764.

Chen,X. et al. (2014) The architecture of a scrambled genome reveals massive levels of genomic rearrangement during development. Cell, 158, 1187–1198.

Cock,P.J.A. et al. (2009) Biopython: freely available Python tools for computational molecular biology and bioinformatics. Bioinformatics, 25, 1422–1423.

Denby Wilkes,C. et al. (2016) ParTIES: a toolbox for Paramecium interspersed DNA elimination studies. Bioinformatics, 32, 599–601.

Feng,Y. et al. (2020) SIGAR: Inferring Features of Genome Architecture and DNA Rearrangements by Split-Read Mapping. Genome Biol. Evol., 12, 1711–1718.

Grüning,B. et al. (2018) Bioconda: sustainable and comprehensive software distribution for the life sciences. Nat. Methods, 15, 475–476.

Guérin,F. et al. (2017) Flow cytometry sorting of nuclei enables the first global characterization of Paramecium germline DNA and transposable elements. BMC Genomics, 18, 327.

Hamilton,E.P. et al. (2016) Structure of the germline genome of Tetrahymena thermophila and relationship to the massively rearranged somatic genome. elife, 5.

Klobutcher,L.A. and Herrick,G. (1995) Consensus inverted terminal repeat sequence of Paramecium IESs: resemblance to termini of Tc1-related and Euplotes Tec transposons. Nucleic Acids Res., 23, 2006–2013.

Li,H. et al. (2009) The Sequence Alignment/Map format and SAMtools. Bioinformatics, 25, 2078–2079.

Ono,Y. et al. (2020) PBSIM2: a simulator for long read sequencers with a novel generative model of quality scores. Bioinformatics.

Sheng,Y. et al. (2020) The completed macronuclear genome of a model ciliate Tetrahymena thermophila and its application in genome scrambling and copy number analyses. Sci. China Life Sci., 63, 1534–1542.

Swart,E.C. et al. (2014) Genome-wide analysis of genetic and epigenetic control of programmed DNA deletion. Nucleic Acids Res., 42, 8970–8983.

Vaser,R. et al. (2017) Fast and accurate de novo genome assembly from long uncorrected reads. Genome Res., 27, 737–746.

Zheng,W. et al. (2020) ADFinder: accurate detection of programmed DNA elimination using NGS high-throughput sequencing data. Bioinformatics, 36, 3632–3636.

