## Supplementary Information for "BleTIES: Annotation of natural genome editing in ciliates using long read sequencing"

Supplements the following article:
"BleTIES: Annotation of natural genome editing in ciliates using long read sequencing"

Brandon K. B. Seah, Estienne C. Swart

### Supplementary Results and Discussion

#### *Benchmarking of BleTIES with simulated long read data*

We simulated long read sequencing data to test BleTIES against a known, defined set of IESs, and to test the effect of different ratios of reads with and without IESs in the library. The genome assembly used for simulation was the relatively well-characterized MAC+IES reference assembly of *Paramecium tetraurelia*, which was derived from experimental knockdown of the PiggyMac gene that resulted in widespread IES retention during development (Arnaiz *et al.*, 2012). Predicted IESs in *Paramecium* are typically bounded by TA repeats, are relatively short (mostly <100 bp), and have a periodic length distribution with peaks every ~10 bp.

A subset of long scaffolds (≥100 kbp) without long gaps (<100 "N" characters) from the MAC+IES assembly and their corresponding MAC contigs was used for read simulation, containing 12199 IES annotations. Although the published *P. tetraurelia* MIC assembly represents a greater fraction of the MIC genome content (Guérin *et al.*, 2017), it is considerably more fragmented (N50 of 37 kbp vs. 431 kbp for the MAC+IES assembly) and was hence less suitable for long read simulation. PacBio subreads and Nanopore reads were simulated from both the MAC+IES and MAC reference assemblies, and mixed at different coverage ratios to represent typical IES retention experiments where only a fraction of reads contain IESs.

##### Comparison of reconstructed to original IESs

As MAC+IES average coverage was increased, the number of predicted IESs increased. As expected, no IESs were predicted at 0× MAC+IES coverage. There was a marked jump in predicted IESs between 10× and 20× (Table S1, Figure S2). Increasing the coverage further to 40× had a much smaller effect. The periodicity observed in IES length distribution was also recovered (Figure S3). At 20× coverage of PacBio subreads, a total of 11312 IESs were predicted (93% of the original 12199), of which 10034 (82%) were at the exact coordinate as the original IES, and a further 1152 (9.4%) within 5 bp of the original IES. Most predicted IESs (9897, 81%) had an exact match to the original coordinate and an assembled sequence flanked by TA repeats. About half (6087, 50%) had an exact matching coordinate, were flanked by TA repeats, and also had the same length as the original IES. At 20× coverage, about half (5845, 48%) of IESs were reconstructed with 100% identity to the original at the correct coordinate, and over two-thirds (8483, 70%) with ≤ 1 mismatch and ≤ 1 indel (Table S2). Simulated Nanopore reads yielded a similar total number of reconstructed IESs, but the IES sequences appeared to be more accurate, with a higher fraction of IESs having the correct length and higher average sequence identity to the original (Tables S1, S2, Figure S2).

### *Benchmarking of BleTIES with real long read data*

Next, we tested BleTIES against real long read data (both PacBio and Nanopore) from *Tetrahymena thermophila*, for which a complete MAC reference assembly (Sheng *et al.*, 2020) as well as a draft MIC assembly (Hamilton *et al.*, 2016) are available. Unlike the simulated *Paramecium* data, the exact number of IESs in the genome is unknown but estimated to be about 12000, as only a curated subset of the IESs have been published (Hamilton *et al.*, 2016). The coverage of the IES-containing reads in the library was also not known *a priori*. *Tetrahymena* IESs are longer on average than those of *Paramecium* (~2000 bp vs. <100 bp), are not flanked by TA repeats, and have imprecise excision, so an IES junction position can vary by several base pairs in a MAC genome library.

#### Comparison of IES reconstructions from PacBio vs. Nanopore reads

A total of 8459 and 8237 IESs were predicted from PacBio and Nanopore libraries respectively, compared to 7510 in the published set (Table S3). Median lengths were similar (1969 vs. 1996 bp respectively) between libraries, and the PacBio library had a higher median retention score (0.48 vs. 0.40) and median (sub)read coverage (63× vs. 52×). A retention score cutoff of 0.1 was chosen after inspection of retention score and IES length histograms, to exclude short, low-scoring IESs (Figure S4), yielding 7930 and 7928 IESs respectively after filtering.

The majority of IES predictions (7574) agree between the PacBio and Nanopore libraries, with coordinates within 50 bp of each other (99% of these were within 11 bp). The median sequence identity was 98.4%, although the PacBio IESs were on average longer than the corresponding Nanopore reconstructions (median +17 bp) (Figure S5).

#### Comparison of reconstructed to published IESs

About two thirds of the predicted IES junctions matched a published IES coordinate within 10 bp, with similar numbers for both the PacBio and Nanopore IESs (Table S3). This left about two thousand previously annotated IESs per library that were not predicted by BleTIES.

IESs in *Tetrahymena* are much longer than those currently predicted in *Paramecium*: the median length of the published IESs was 2775 bp (max 43 kbp), whereas most *Paramecium* IESs were < 100 bp. Therefore, we hypothesize that BleTIES did not predict many IESs because the reads were not long enough to completely span them. Published IESs that were not predicted by BleTIES were generally longer than those that were predicted (median 5345 vs 2114 bp for PacBio, 5600 vs. 2100 bp for Nanopore) (Figure S6). When we inspected read mappings to the MAC reference for several published IESs that were not predicted by BleTIES, we found that the IES-containing portion of the read was usually reported by the mapper as a clip, rather than an insert, consistent with our hypothesis.

BleTIES also predicted new IESs that were not previously annotated (Table S3). These have similar lengths and retention scores to the other BleTIES predicted IESs, although there is a small fraction with low retention scores and lengths that probably represent spurious predictions despite the earlier filtering, or potentially also “cryptic” IESs that are excised at lower frequency than regular IESs (Duret *et al.*, 2008). Inspection of read mappings for several newly predicted IESs showed clusters of sequence inserts of similar length and sequence around the mapping position, and with no obvious signs of mismapping, suggesting that these are mostly authentic IESs. When the BleTIES and published IESs are pooled, there are about 10000 IES annotations, which approaches an earlier estimate of 12000 for the actual number of IESs in the *Tetrahymena* MIC genome.

For IESs that overlapped between the predicted and published sets, BleTIES IES reconstructions had high identity to the published sequences (median 99.0% and 98.8% for PacBio and Nanopore respectively). IES reconstructions from PacBio reads were on average slightly longer than the published sequence (median +9 bp) whereas those from Nanopore were slightly shorter (median -8 bp) (Figure S7).

If metadata linking PacBio subreads to their original zero mode waveguide (ZMW) are available, the actual physical coverage of each IES could be reported separately from the subread coverage, because each ZMW represents a single DNA molecule, from which multiple subreads can be produced. This information is parsed by BleTIES MILRAA from the subread names in the read file. Regrettably, subread headers are typically renamed when the sequences are deposited in the NCBI Short Read Archive, removing the original metadata, so the physical coverage could not be recovered in this case.

### Supplementary Methods

#### *Simulation of PacBio subreads and Nanopore reads from* Paramecium

The MAC+IES and MAC reference assemblies (both v1.0) for *Paramecium tetraurelia* strain 51 were downloaded from ParameciumDB https://paramecium.i2bc.paris-saclay.fr (Arnaiz *et al.*, 2020). MAC+IES scaffolds containing ≥100 N's and/or with length ≤ 100 kbp were removed; their corresponding MAC scaffolds were also removed. The remaining scaffolds, containing 12119 IESs, were used to simulate long reads with pbsim2 v2.0.1 https://github.com/yukiteruono/pbsim2 (Ono *et al.*, 2020), using either the provided PacBio P6-C4 sequencing chemistry or Oxford Nanopore R9.5 error models, random seed --seed 12345, and default length parameters: --length-min 100 --length-max 1000000 --length-mean 9000 --length-sd 7000. The difference ratio parameter was set to default 6:50:54 for PacBio, and 23:31:46 for Nanopore, as recommended in the pbsim2 documentation. For each sequencing platform, separate simulated libraries were produced for MAC+IES and MAC, to an average coverage of 50x each. The MAC+IES and MAC libraries were then subsampled with seqtk v1.3-r116-dirty https://github.com/lh3/seqtk to generate mixed libraries in the following average coverage ratios: 0:50, 5:45, 10:40, 20:30, 40:10.

#### *Mapping and IES prediction from* Paramecium *simulated reads*

The mixed MAC+IES : MAC simulated libraries were mapped against the MAC reference assembly with minimap2 v2.17-r974-dirty https://github.com/lh3/minimap2 (Li, 2018), using the -ax map-pb option for PacBio and -ax map-ont for Nanopore. The mapped reads were converted to BAM, sorted, and indexed with samtools v1.9 (Li *et al.*, 2009). IESs were predicted from each mapped library with BleTIES MILRAA v0.1.9 with options --type subreads --min_break_coverage 5 --min_del_coverage 5.

#### *Data sources for* Tetrahymena *long reads and reference genomes*

PacBio Sequel I subreads and Nanopore MinION reads from a *Tetrahymena thermophila* MIC enrichment sample (SAMN12736122) under project accession PRJNA565159 were downloaded from the NCBI Short Read Archive. Reference genome assemblies and feature tables for MAC (2021 version) (Sheng *et al.*, 2020) and MIC (2016 version) (Hamilton *et al.*, 2016) were downloaded from the Tetrahymena Genome Database (http://ciliate.org/index.php/home/downloads) (Stover *et al.*, 2012).

Coordinates of IES junctions relative to the MAC reference genome were not available. To obtain these, we first extracted the 500 bp flanking each IES annotated in the MIC assembly, and concatenated this into a 1 kbp sequence, which was then aligned with BLAT v36 (Kent, 2002) to the MAC assembly. Because IESs are imprecisely excised in *Tetrahymena*, these were frequently aligned in two blocks of approximately 500 bp each. Flanking sequences that aligned in one or two blocks, or comprising three or four blocks where one block was >480 bp, were used to identify IES junctions and orientation relative to the MAC reference. Of 7544 IESs annotated in the MIC, corresponding MAC coordinates could be found for 7510.

#### *Mapping and IES prediction from* Tetrahymena *long reads*

PacBio and Nanopore reads were separately mapped to the MAC reference genome, with the same parameters as simulated reads, described above. For each sequencing platform, mappings were merged, sorted, and indexed with samtools into a single BAM file. IESs were predicted for each sequencing platform with BleTIES MILRAA as described above. Only IESs with retention scores ≥0.1 were used for downstream comparisons.

#### *Comparison of reconstructed to annotated IES sequences*

For simulated data from *Paramecium*, BleTIES IES predictions were matched to existing published IES annotations if the coordinate was exactly the same (at the TA boundary if present). For *Tetrahymena thermophila*, they were matched if within 10 bp from each other, because IESs in that species are imprecisely excised (i.e. the excision boundaries can vary by a few bp). BleTIES and published IES sequences were aligned with the function pairwise2.align.globalxx from Biopython v1.76 (Cock *et al.*, 2009), returning the alignment score (number of matching bases). Sequence identity was calculated as the ratio of alignment score to the longer of the two lengths, number of mismatches as the difference between published IES length and alignment score, and net indels as the length difference between BleTIES IES sequence and published sequence.

#### *Code availability*

Scripts and computational notebooks documenting the above tests are available at https://github.com/Swart-lab/bleties-test-ptet and https://github.com/Swart-lab/bleties-test-tthe.

### Supplementary References

Bibliography


Arnaiz,O. *et al.* (2020) ParameciumDB 2019: integrating genomic data across the genus for functional and evolutionary biology. *Nucleic Acids Res.*, **48**, D599–D605.

Arnaiz,O. *et al.* (2012) The Paramecium germline genome provides a niche for intragenic parasitic DNA: evolutionary dynamics of internal eliminated sequences. *PLoS Genet.*, **8**, e1002984.

Cock,P.J.A. *et al.* (2009) Biopython: freely available Python tools for computational molecular biology and bioinformatics. *Bioinformatics*, **25**, 1422–1423.

Duret,L. *et al.* (2008) Analysis of sequence variability in the macronuclear DNA of Paramecium tetraurelia: a somatic view of the germline. *Genome Res.*, **18**, 585–596.

Guérin,F. *et al.* (2017) Flow cytometry sorting of nuclei enables the first global characterization of Paramecium germline DNA and transposable elements. *BMC Genomics*, **18**, 327.

Hamilton,E.P. *et al.* (2016) Structure of the germline genome of Tetrahymena thermophila and relationship to the massively rearranged somatic genome. *elife*, **5**.

Kent,W.J. (2002) BLAT--the BLAST-like alignment tool. *Genome Res.*, **12**, 656–664.

Li,H. (2018) Minimap2: pairwise alignment for nucleotide sequences. *Bioinformatics*, **34**, 3094–3100.

Li,H. *et al.* (2009) The Sequence Alignment/Map format and SAMtools. *Bioinformatics*, **25**, 2078–2079.

Ono,Y. *et al.* (2020) PBSIM2: a simulator for long read sequencers with a novel generative model of quality scores. *Bioinformatics*.

Sheng,Y. *et al.* (2020) The completed macronuclear genome of a model ciliate Tetrahymena thermophila and its application in genome scrambling and copy number analyses. *Sci. China Life Sci.*, **63**, 1534–1542.

Stover,N.A. *et al.* (2012) Tetrahymena Genome Database Wiki: a community-maintained model organism database. *Database (Oxford)*, **2012**, bas007.

### Supplementary Tables

*Table S1*. Comparison of reconstructed IESs from simulated PacBio (PB) or Nanopore (ONT) reads vs. original MAC+IES annotations, for increasing average coverages of the MAC+IES component in simulated read libraries.

| | **PacBio** | | | | **Nanopore** | | | |
|---|---|---|---|---|---|---|---|---|
| **MAC+IES average coverage** | **5** | **10** | **20** | **40** | **5** | **10** | **20** | **40** |
| **Coordinate within 10 bp** | 4 | 49 | 96 | 98 | 2 | 43 | 66 | 67 |
| **Coordinate within 5 bp** | 37 | 675 | 1152 | 1015 | 29 | 439 | 805 | 735 |
| **Coordinate match** | 4 | 92 | 127 | 115 | 1 | 41 | 53 | 42 |
| **Coordinate match, TA-flanked** | 91 | 2184 | 3810 | 3492 | 69 | 1245 | 1679 | 1545 |
| **Coordinate & length match, TA-flanked** | 133 | 2498 | 6087 | 6886 | 207 | 3726 | 8645 | 9188 |
| **Coordinate & length match** | 0 | 4 | 10 | 6 | 0 | 8 | 6 | 1 |
| **No match** | 0 | 10 | 30 | 30 | 0 | 13 | 22 | 31 |
| **Total** | 269 | 5512 | 11312 | 11642 | 308 | 5515 | 11276 | 11609 |

*Table S2*. Comparison of sequence identity, number of mismatches, and number of indels between original and reconstructed IESs from simulated PacBio subreads (PB) and Nanopore reads (ONT), as the average coverage of MAC+IES in simulated read libraries was varied.

| | **PacBio** | | | | **Nanopore** | | | |
|---|---|---|---|---|---|---|---|---|
| **MAC+IES average coverage** | **5** | **10** | **20** | **40** | **5** | **10** | **20** | **40** |
| **100% identity** | 126 | 2367 | 5845 | 6679 | 193 | 3535 | 8425 | 8975 |
| **≥97% identity** | 191 | 4011 | 8931 | 9545 | 264 | 4730 | 10058 | 10503 |
| **0 mismatch** | 206 | 4228 | 9140 | 9681 | 225 | 4192 | 9323 | 9781 |
| **≤1 mismatch** | 223 | 4678 | 9911 | 10357 | 267 | 4835 | 10159 | 10601 |
| **0 indel** | 133 | 2502 | 6097 | 6892 | 207 | 3734 | 8651 | 9189 |
| **≤1 indel** | 183 | 3837 | 8541 | 9099 | 265 | 4766 | 10074 | 10486 |
| **≤1 mismatch and ≤1 indel** | 181 | 3791 | 8483 | 9041 | 260 | 4680 | 9976 | 10414 |
| **Total with matching coordinate** | 228 | 4778 | 10034 | 10499 | 277 | 5020 | 10383 | 10776 |

*Table S3.* Metrics of IES predictions from real *Tetrahymena thermophila* long read data, and their overlap with previously annotated IESs from the MIC reference assembly.

| | **PacBio** | **Nanopore** |
|---|---|---|
| **Total IESs predicted** | 8459 | 8237 |
| **Retention score > 0.1** | 7930 | 7928 |
| **Published IESs within 10 bp of predicted** | 5128 | 5087 |
| **Published IESs with no MILRAA IESs within 10 bp** | 2382 | 2431 |
| **MILRAA IESs with no published IESs within 10 bp** | 2825 | 2872 |

### Supplementary Figures

*Figure S1*. Flowchart of BleTIES pipeline showing required input data (yellow), BleTIES modules (pink), and pipeline outputs (blue).

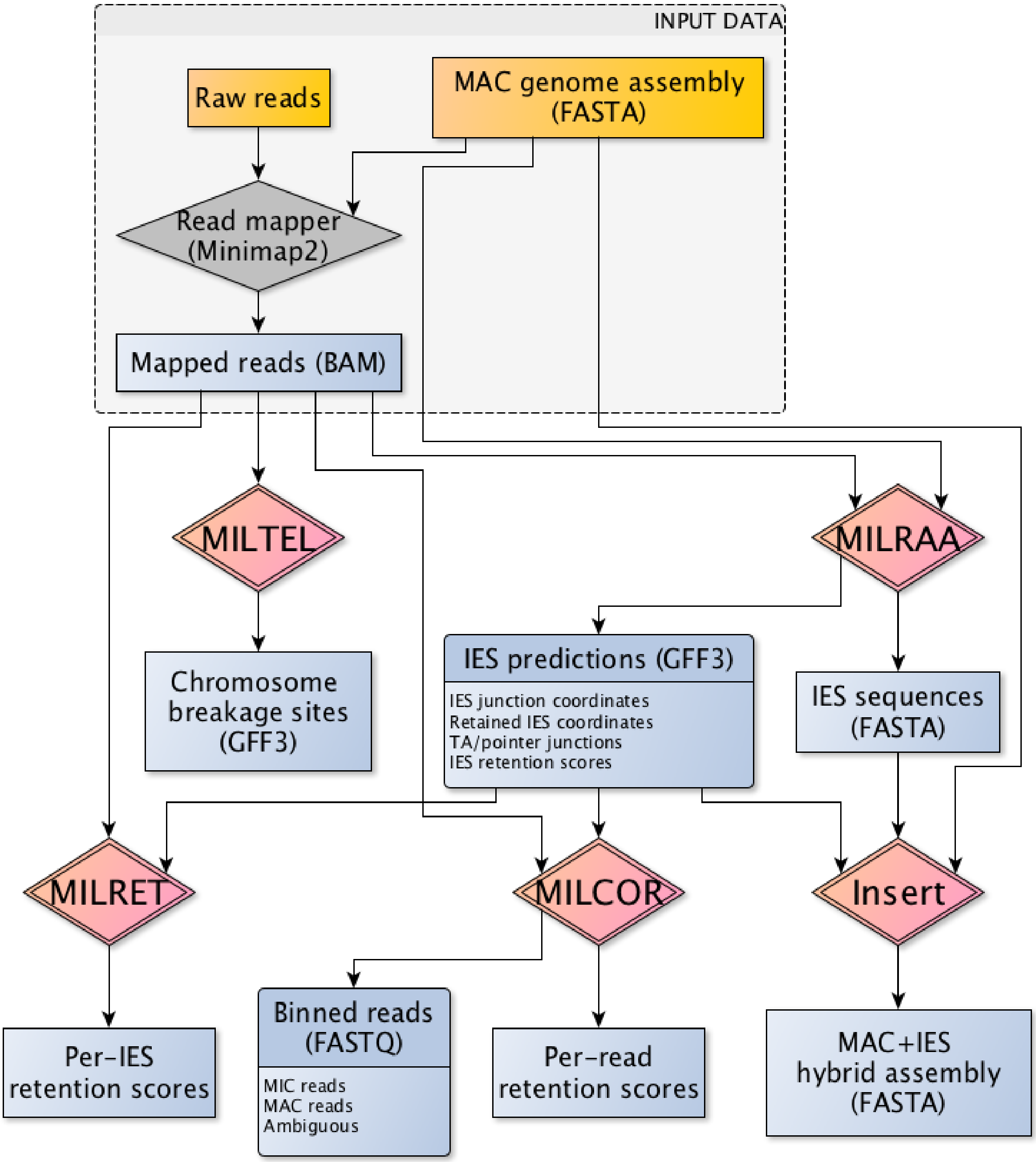

*Figure S2*. Barplots of total numbers of predicted IESs (*vertical axis*) from simulated libraries containing mixtures of MAC+IES and MAC reads, as the average coverage of the MAC+IES component was varied (*horizontal axis*), for PacBio subreads (*above*) and Nanopore (*below*) platforms. Each bar is further broken down into how well the reconstructed sequence matched the original IES sequence, in terms of insert coordinate (coord), presence of flanking TAs (TAs), and sequence length.

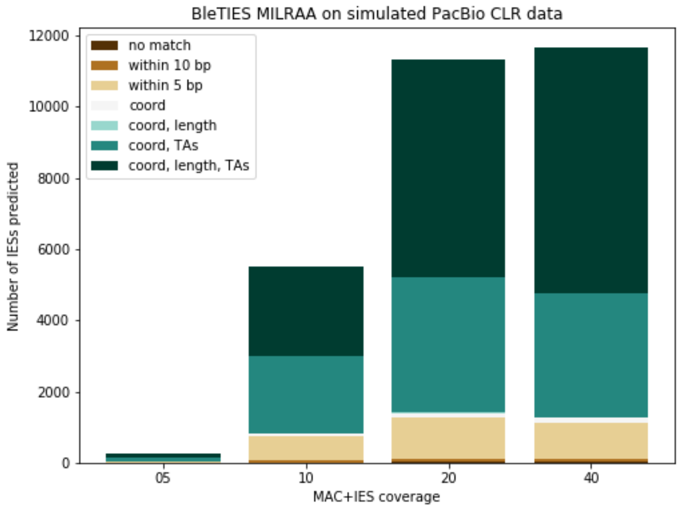


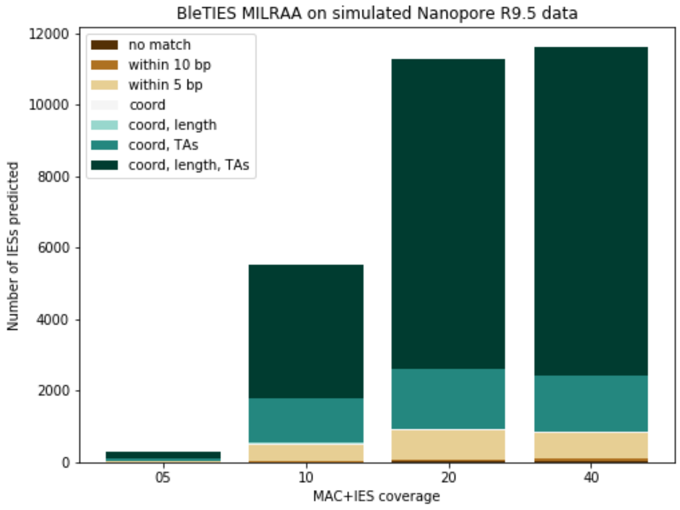

*Figure S3.* Histograms of reconstructed IES lengths from simulated PacBio (*left column*) and Nanopore (*right column*) libraries, as average coverage of MAC+IES reads was varied (rows), showing only lengths ≤ 300 bp.

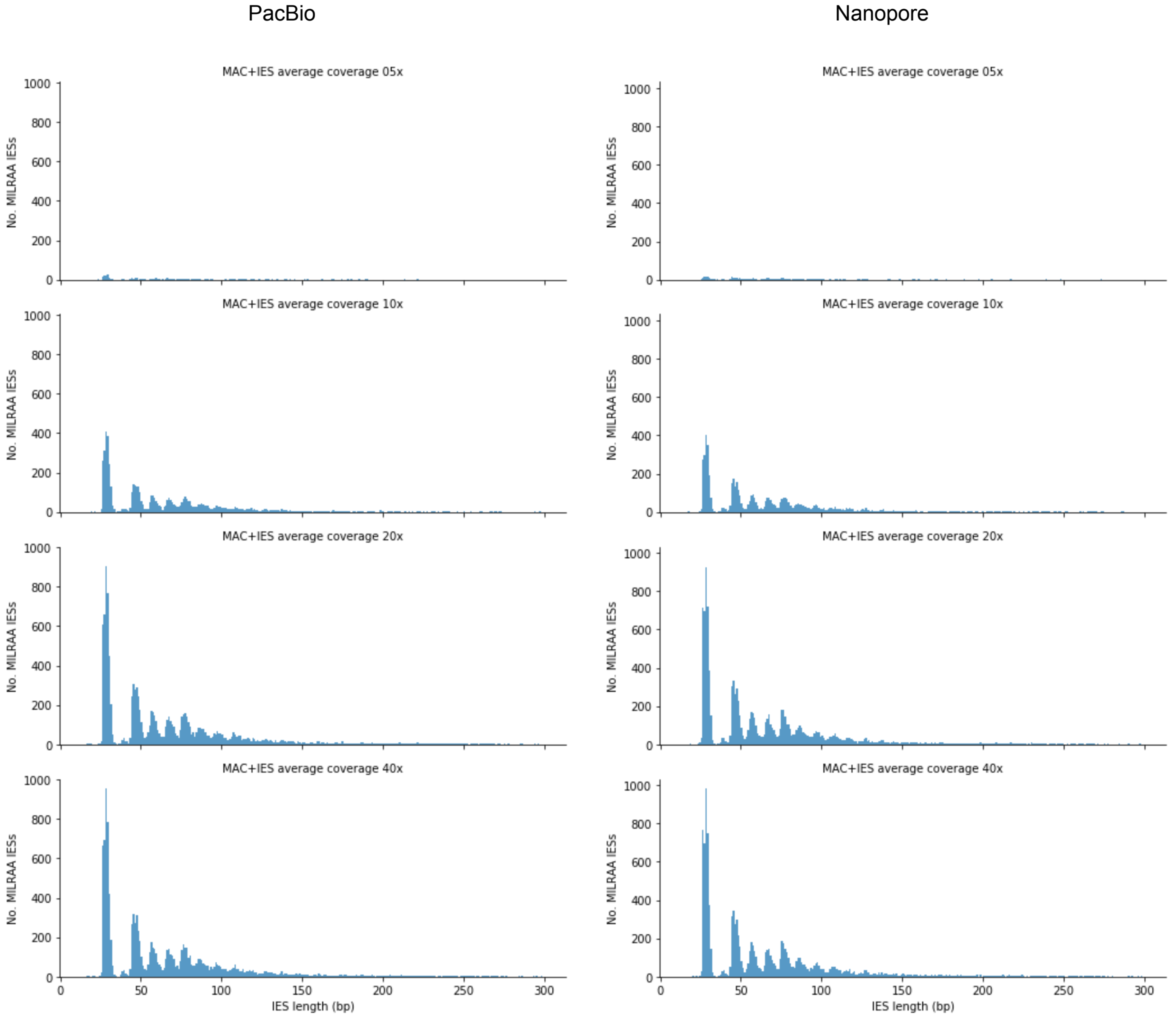

*Figure S4*. Histograms of reconstructed IES lengths (*vertical axis*) vs. retention scores (*horizontal axis*) from PacBio (*left*) vs. Nanopore (*right*) libraries for *Tetrahymena thermophila*, suggesting a retention score cutoff of 0.1 to remove low-scoring and likely spurious predictions.

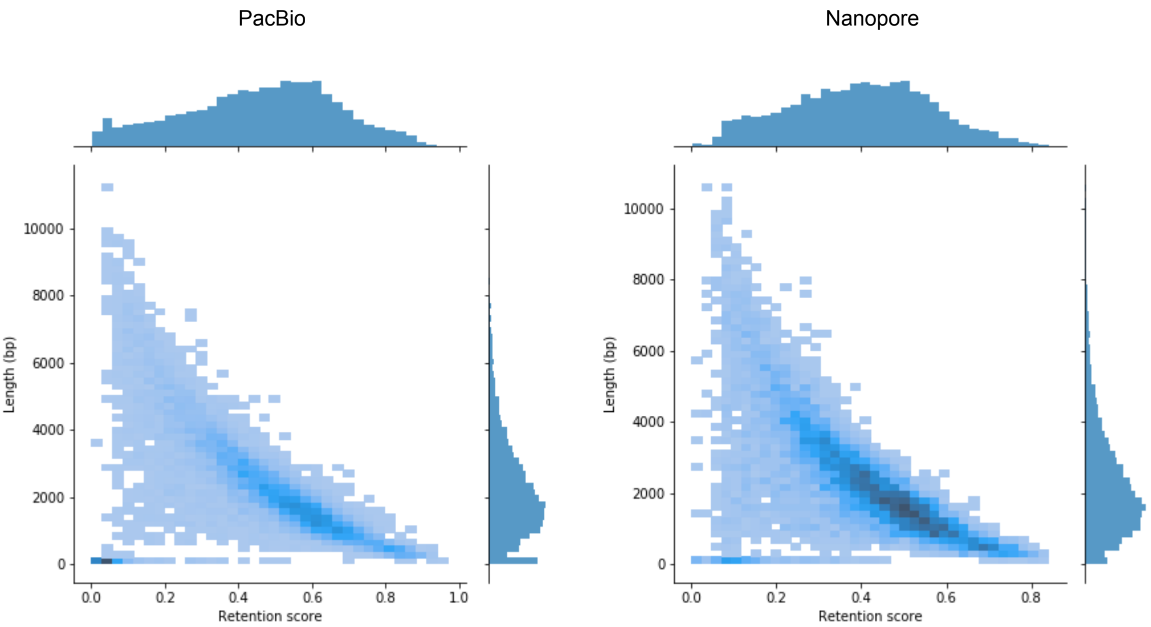

*Figure S5*. Comparison of *Tetrahymena* IES sequences predicted from PacBio vs. from Nanopore read libraries in terms of sequence identity (*upper left*), number of mismatched bases (*upper right*), length difference relative to the Nanopore version (*lower left*), and difference in the predicted insert position on the reference MAC genome (*lower right*).

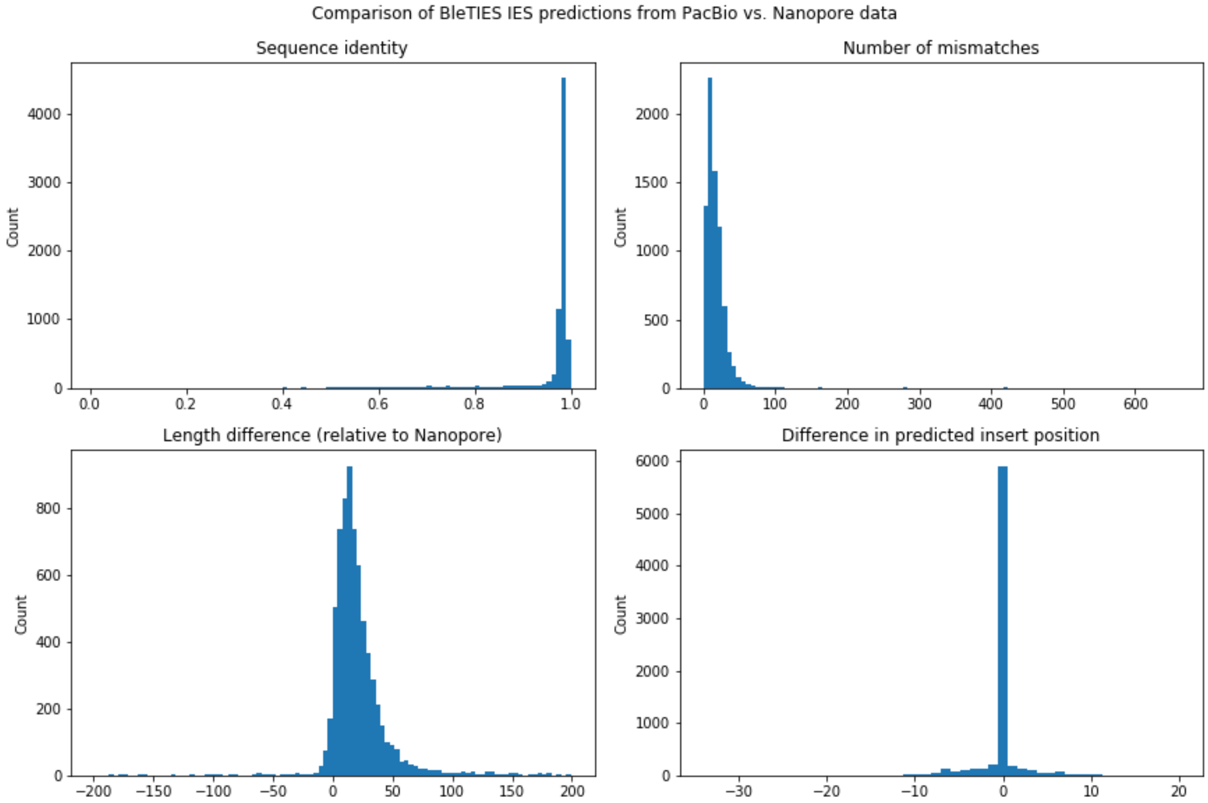

*Figure S6*. Lengths of previously annotated *Tetrahymena* IESs, comparing those that were also predicted by BleTIES (*orange*) vs. those that were not (*blue*), from either PacBio read library (*left*) or Nanopore read library (*right*).

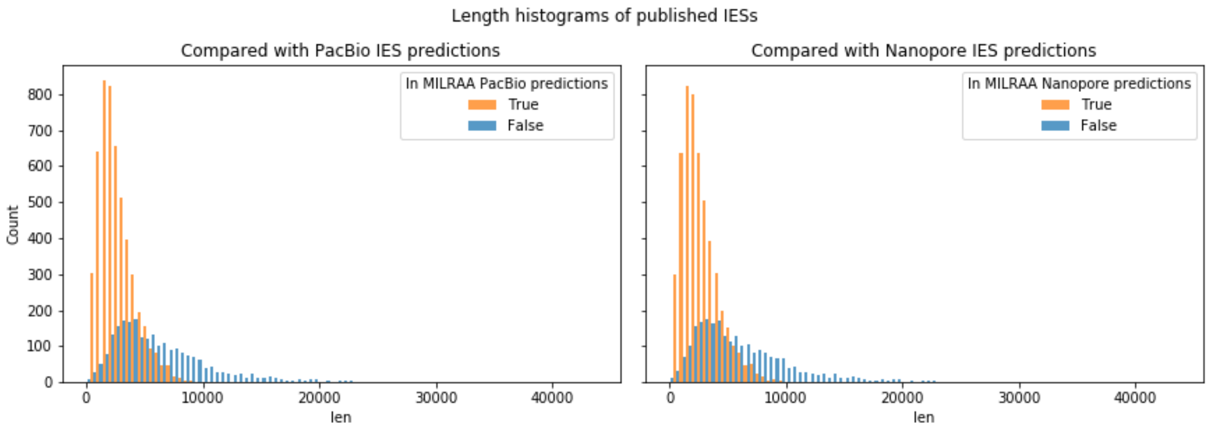

*Figure S7*. Comparison of reconstructed *Tetrahymena* IES sequences from PacBio (*left column*) or Nanopore (*right column*) read libraries to the previously annotated IES sequences, in terms of sequence identity (*top row*), number of mismatches to the published sequence (*middle row*), and length difference relative to the published sequence (*bottom row*).

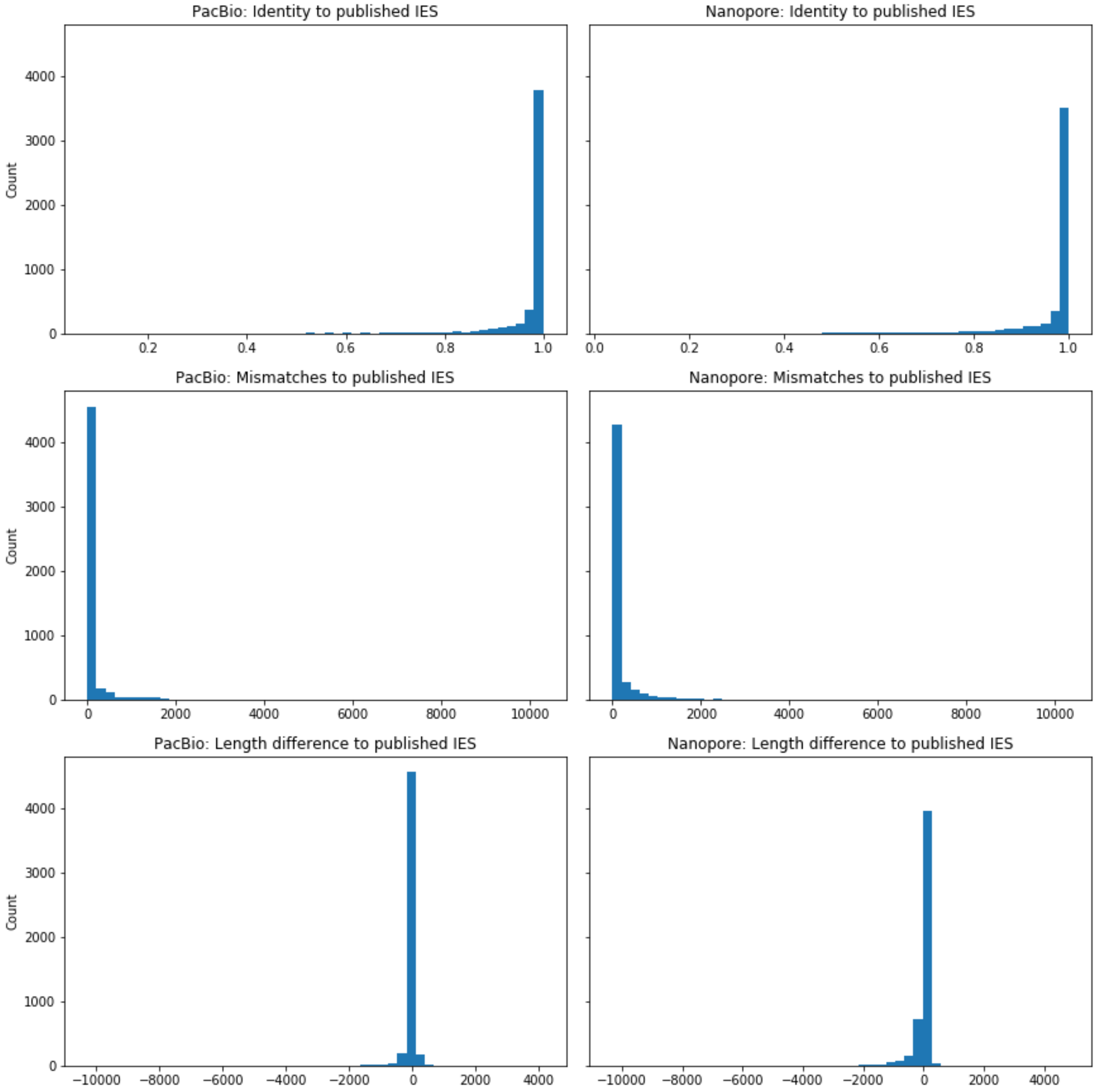
